# Human pre-60S assembly factors link rRNA transcription to pre-rRNA processing

**DOI:** 10.1101/2022.03.01.482553

**Authors:** Amber F. Buhagiar, Mason A. McCool, Carson J. Bryant, Laura Abriola, Yulia V. Surovtseva, Susan J. Baserga

## Abstract

In eukaryotes, the nucleolus is the site of ribosome biosynthesis, an essential process in all cells. While human ribosome assembly is largely evolutionarily conserved, many of the regulatory details underlying its control and function have not yet been well-defined. The nucleolar protein RSL24D1 was originally identified as a factor important for ribosome biogenesis, and as an interactor with the PeBoW complex (PES1, BOP1, WDR12) in high-throughput affinity purifications. The PeBoW complex has been shown to be required for pre-28S rRNA processing. In this study, we show that RSL24D1 depletion impairs both pre-ribosomal RNA (pre-rRNA) transcription and mature 28S rRNA production, leading to decreased protein synthesis and p53 stabilization in mammalian cells. Surprisingly, each of the PeBoW complex members is also required for pre-rRNA transcription. We also demonstrate that RSL24D1 is physically complexed with RNA polymerase I, revealing a connection between large ribosomal subunit biogenesis and rDNA transcription. These results uncover the dual role of RSL24D1 and the PeBoW complex in multiple steps of ribosome biogenesis, and provide evidence implicating large subunit biogenesis factors in pre-rRNA transcription control.

## Introduction

Ribosome biogenesis (Ribi) is a highly conserved and essential process and among eukaryotes has been most thoroughly investigated in baker’s yeast, *Saccharomyces cerevisiae* (*S. cerevisiae*) (Woolford and Baserga 2013). The two ribosomal subunits, large 60S (LSU) and small 40S (SSU), are assembled independently in the nucleolus and join in the cytoplasm to translate mRNAs to proteins. The process begins with RNA polymerase I (RNAPI) driving transcription of the ribosomal DNA (rDNA) to produce the 47S pre-rRNA transcript in humans. The transcript is subsequently processed and modified by a wide range of transacting factors to produce the 18S, 5.8S, and 28S mature rRNAs that form the SSU (18S) and LSU (5.8S, 28S, and RNAPIII-transcribed 5S) respectively. As rDNA transcription is the rate-limiting step of ribosome biosynthesis which underlies cell growth and proliferation, this process is under tight regulation to ensure cellular homeostasis.

Pre-rRNA processing and ribosome assembly require the hierarchical association and dissociation of a large number of assembly factors. Some of these factors are required for the formation of both ribosomal subunits, while others are required for the synthesis of one of the two subunits. Moreover, a group of factors known collectively as the transcription U3 small nucleolar RNA Associated Proteins (t-UTPs), a subcomplex of the SSU processome, are required for the early pre-rRNA transcription steps in eukaryotic cells in addition to their functional role in the processing of the 18S rRNA (Gallagher et al. 2004; Krogan et al. 2004; Prieto and McStay 2007). The t-UTPs associate with the SSU processome to coordinate this action (Krogan et al. 2004). While SSU processome components exhibit dual-functional roles in transcription and processing, it remains largely unknown whether there are nucleolar ribosome assembly factors required for LSU biogenesis that play a similar role in RNAPI transcription.

RSL24D1 is an evolutionarily conserved protein previously studied in yeast. The *S. cerevisiae* Rlp24 protein (ortholog of mammalian RSL24D1) is required for 27SB pre-rRNA processing to 5.8S and 25S rRNAs via ITS2 cleavage to produce the 60S ribosomal subunit (Kappel et al. 2012). Rlp24 associates with both early and late stage pre-60S particles in the nucleolus and cytoplasm respectively, and is replaced by the homologous ribosomal protein eL24 via the AAA-ATPase Drg1 (Kappel et al. 2012). In humans, RSL24D1 expression in tumor educated platelets has been negatively correlated with early-stage cancer progression and its upregulation has been associated with familial hypercholesterolemia (Li et al. 2015; Ge et al. 2021). Our laboratory’s previously conducted RNAi screening campaign identified RSL24D1 as a factor important for the synthesis of ribosomes in mammalian cells (Farley-Barnes et al. 2018). However, the molecular mechanisms of how RSL24D1 participates in ribosome assembly in metazoan cells have not yet been fully understood.

In this study we establish mammalian RSL24D1 as a critical factor for LSU production. It is required for normal nucleolar number in MCF10A cells, a highly-predictive indicator of a function in Ribi (Farley-Barnes et al. 2018; Ogawa et al. 2021). More specifically, we confirm that RSL24D1 is required for cleavage at site 2 in ITS2 of pre-rRNAs to produce the 60S large subunit. As a pre-rRNA processing factor, we also show that RSL24D1 interacts with members of the PeBoW (PES1-BOP1-WDR12) complex. Unexpectedly, we uncover a previously unidentified role for RSL24D1 and PeBoW in RNAPI transcription. These results support the critical role for RSL24D1 in rRNA synthesis and reveal a putative link between LSU biogenesis and RNAPI transcription regulation.

## Results

### RSL24D1 is required to maintain nucleolar number and viability in MCF10A human breast epithelial cells

Previously, we performed a genome-wide, phenotypic RNAi screen to uncover novel protein regulators of nucleolar number using a library of siGENOME siRNAs (Horizon Discovery) (Farley-Barnes et al. 2018; Ogawa et al. 2021). The screen was performed in human breast epithelial MCF10A cells, a near-diploid non-cancerous cell line, which contain an average of 2-3 nucleoli per cell nucleus. Proteins whose depletion led to a significant change in nucleolar number, either a decrease to 1 or an increase to 5 or more, were identified as candidate novel regulators of ribosome biogenesis. RSL24D1 depletion was observed to cause a striking decrease in nucleolar number relative to the siGFP negative control and siUTP4 positive control (Fig. 1A). To minimize the inclusion of off-target effect induced observations, we rescreened RSL24D1 using ON-TARGETplus siRNA reagents (siONT; Horizon Discovery; Data S1), which are designed to reduce off-target effects. We found that RSL24D1 depletion using a siONT pool reproduces the one-nucleolus phenotype relative to the non-targeting (siNT) negative control and the siNOL11 positive control (Fig. 1B; Table S1).

**Figure 1:**
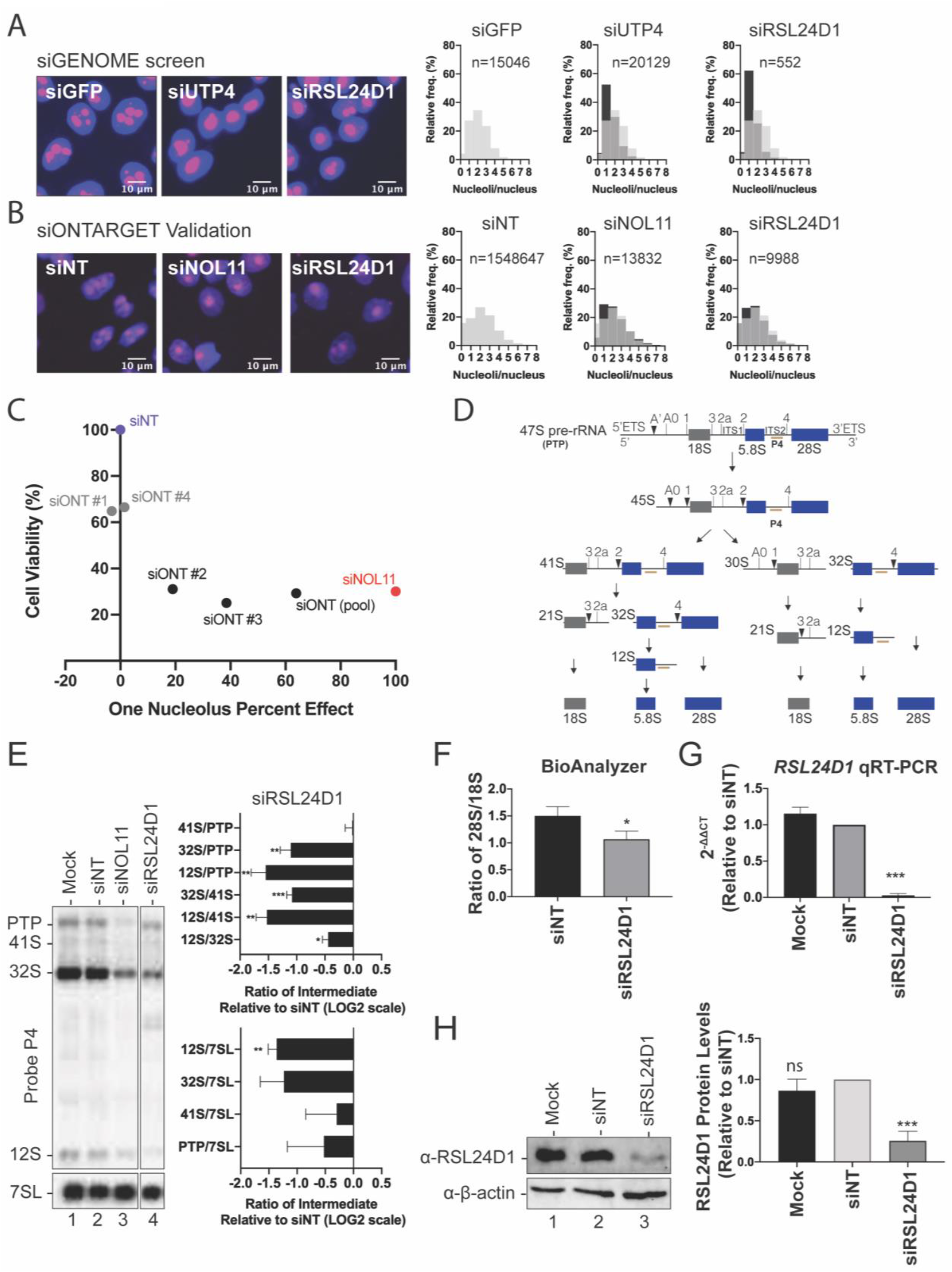
RSL24D1 functions in large subunit pre-rRNA processing. **(A, B)** RSL24D1 siRNA depletion causes a decrease in nucleolar number in MCF10A cells. **(A)** RSL24D1 depletion using siGENOME siRNAs decreases nucleolar number. Left panel: Representative images of nuclei stained with Hoechst (blue) and nucleoli stained with an anti-fibrillarin antibody (pink) siGFP was used as a negative control (2-3 nucleoli/nucleus) and siUTP4 was used as a positive control (1 nucleolus/nucleus). Right panel: Histograms of the relative frequency of nucleoli per nucleus are shown for the controls and siRSL24D1. There is a decrease in nuclei with 2-3 nucleoli for positive controls and RSL24D1-depleted cells. Light gray bars show the nucleolar number distribution for siGFP negative control, and black bars show the nucleolar number distribution for indicated positive control or siRSL24D1, with number of cells analyzed for each treatment indicated. **(B)** RSL24D1 depletion using ON-TARGETplus siRNA reagents decreases nucleolar number. Panels as above, except siNT was used as a negative control and siNOL11 was used as a positive control. **(C)** Two of four RSL24D1 siRNAs (#2 and #3) decrease both nucleolar number and cell viability in MCF10A cells. Oligonucleotide deconvolution of the ON-TARGETplus siRNA pool was used to confirm the activity of individual siRNAs targeting RSL24D1 in the assay for nucleolar number and for cell viability. Key: *RSL24D1* targeting siONTs that decreased nucleolar number (black), *RSL24D1* targeting siONTs that did not decrease nucleolar number (gray), siNT negative control (blue), and siNOL11 positive control (red). **(D)** Mammalian pre-rRNA processing schematic. The probe P4 hybridizes with the ITS2 region of the 47S pre-rRNA precursor which undergoes a series of cleavage steps to yield the small subunit 18S rRNA (gray) and the large subunit 5.8S and 28S rRNAs (blue). **(E)** RSL24D1 is required for ITS2 processing. Representative northern blots measuring steady-state levels of pre-rRNA intermediates when RSL24D1 is depleted in MCF10A cells are shown. A probe for the 7SL RNA was used as a loading control. Negative controls were mock (no siRNA) and siNT (non-targeting) and siNOL11 was used as a positive control. Defects in processing were determined by analyzing precursor-product relationships using Ratio Analysis of Multiple Precursors (RAMP) and were reported relative to siNT. Quantitation by RAMP is shown to the right. Graphs indicate mean ± SEM, n = 3 biological replicates. Data were analyzed by two-way ANOVA using GraphPad Prism. **(F)** RSL24D1 depletion in MCF10A cells decreases the 28S/18S mature rRNA ratio by Agilent BioAnalyzer. Graph indicates mean ± SD, n = 3 biological replicates. Data were analyzed by Student’s t-test in GraphPad Prism where * p ≤ 0.05. **(G)** qRT-PCR confirmation of *RSL24D1* knockdown in MCF10A cells. 2^−ΔΔCt^ values, relative to a siNT control and 7SL internal control primer, show knockdown of *RSL24D1* by qRT-PCR using the indicated siRNAs. Graph indicates mean ± SD, n = 3 biological replicates. Data were analyzed by Student’s t-test followed by multiple testing p-value correction (two-stage linear step-up procedure of Benjamini, Krieger, and Yekutieli) using GraphPad Prism where *** p ≤ 0.005. **(H)** Western blot confirmation of RSL24D1 knockdown in MCF10A cells. Mock and non-targeting (siNT) siRNAs are shown as negative controls. Quantitation of RSL24D1 levels (right panel) is reported relative to siNT and normalized to the β-actin loading control. Graph indicates mean ± SD, n = 3 biological replicates. Data were analyzed by one-way ANOVA with Dunnett’s multiple comparisons test in GraphPad Prism where *** p ≤ 0.001; ns = not significant.

We performed deconvolution of the RSL24D1 siRNA pool to further validate the role of RSL24D1 in maintaining normal nucleolar number. The individual siRNAs from the siONT pool of four siRNAs were evaluated for their effects on nucleolar number and cell viability in RSL24D1-depleted cells. We found that 2 of the 4 individual siRNAs significantly reduced the number of nucleoli from 2-3 to only 1, which also corresponds with significantly reduced cell viability relative to the two siRNAs that did not significantly reduce nucleolar number (Fig. 1C; Table S1; Fig. S1). A correlation between the presence of one nucleolus upon RSL24D1 depletion and cell viability is expected if RSL24D1 is required for ribosome biogenesis, an essential cellular process. Thus, we hypothesized that the reduction in nucleolar number upon RSL24D1 depletion was the result of defective ribosome biogenesis.

Changes in nucleolar number and morphology have been connected with cancer pathology and prognosis for over one-hundred years (Derenzini et al. 2017). Based on our result that RSL24D1 is necessary for maintenance of nucleolar number, we examined *RSL24D1*’s mRNA expression levels in Genotype-Tissue Expression (GTEx) unmatched normal and The Cancer Genome Atlas (TCGA) matched normal and tumor samples (Goldman et al. 2020). Although *RSL24D1* expression is negatively correlated with cancer progression in tumor educated platelets (Ge et al. 2021), this is not what we observed for *RSL24D1*’s expression in tumor compared to normal tissue. *RSL24D1* was significantly higher expressed in all tumor samples, including breast tumor, compared to normal tissue (Fig. S2). *RSL24D1*’s increased expression in breast cancer highlights the relevance of its expected role in ribosome biogenesis in MCF10A cells.

### RSL24D1 functions in the processing of pre-28S rRNA

RSL24D1 was identified in a previous targeted RNAi depletion screen as a factor potentially involved in LSU processing in HeLa cells and 60S biogenesis in mouse embryonic stem cells (Tafforeau et al. 2013; Durand et al. 2021). Moreover, the RSL24D1 yeast ortholog, Rlp24, has been shown to be important for large subunit pre-rRNA processing (Kappel et al. 2012). A key step during large subunit synthesis is the cleavage of ITS2 from maturing pre-60S subunits. In the 47S primary transcript precursor (PTP), ITS2 separates the 5.8S and 28S rRNAs and is eventually removed from the larger pre-rRNA precursor (Fig. 1D). We probed the extent to which pre-rRNA processing was disrupted upon depletion of RSL24D1 using siONT pools in MCF10A cells by northern blot analysis, using the ITS2 probe P4 to detect defects in LSU biogenesis. Mock (no siRNA) and siNT were used as negative controls and siNOL11 was used as a positive control. After 72 h of siRNA knockdown, RSL24D1 depletion resulted in a significant decrease in the 32S and 12S pre-rRNAs relative to the mock and non-targeting siRNA controls (Fig. 1E). These results concur with previous northern blot results where 2 of 3 siRNAs against *RSL24D1* mRNA effected the same reduction in 12S pre-rRNA (Tafforeau et al. 2013). Quantitation of the ratios of each intermediate relative to its precursor in the processing pathway by the Ratio Analysis of Multiple Precursors (RAMP) method (Wang et al. 2014) revealed that the 12S/32S, 12S/41S, and 12/PTP ratios were all significantly decreased after depletion of RSL24D1. Although depletion of NOL11 led to reduced 32S pre-rRNA, NOL11 depletion did not decrease the 12S/32S, 12S/41S, and 12S/PTP ratios, consistent with its role in pre-18S rRNA processing.

Because the pre-rRNA processing defects indicate aberrant LSU biogenesis, we sought to determine the extent to which RSL24D1 depletion by siONT pools affects the production of the mature 28S rRNA. Agilent BioAnalyzer quantitation shows a decrease in the ratio of mature 28S to 18S levels (Fig. 1F) indicating less LSU rRNA (28S) relative to SSU (18S). Furthermore, *RSL24D1* mRNA levels and RSL24D1 protein levels were greatly reduced following RSL24D1 siONT pool depletion, indicating robust knockdown using this method (Fig. 1G, H). We confirm that RSL24D1 is required for pre-LSU rRNA processing and for maintaining levels of the 28S rRNA.

### RSL24D1 interacts with the PeBoW complex proteins in humans

To gain further biological insight into the role of RSL24D1, we employed the search tool for retrieval of interacting genes (STRING) and the human protein complex map (hu.MAP 2.0) to predict functional associations of RSL24D1 (Szklarczyk et al. 2015; Drew et al. 2020). hu.MAP 2.0 incorporates information from several recently published protein interaction and affinity purification mass spectrometry datasets to predict and catalog mammalian protein complexes. Of the medium-to-high confidence interacting proteins identified, WDR12 was shown to interact with RSL24D1 in both datasets (Table 1; Table S2). WDR12 is a member of the PES1-BOP1-WDR12 (PeBoW) nucleolar complex that is essential for large subunit processing and maturation (Rohrmoser et al. 2007). Moreover, BOP1 and PES1 were also among the RSL24D1-interacting proteins revealed by hu.MAP 2.0. Overexpression of PeBoW proteins has been associated with several cancers and is thought to promote aberrant cellular proliferation by stimulating ribosome biogenesis (Killian et al. 2006; Fan et al. 2018; Yin et al. 2018). Interestingly, the yeast complex orthologous to PeBoW (Nop7 complex) containing factors Nop7, Erb1, and Ytm1 has been shown to interact with Rlp24, the yeast ortholog of RSL24D1 (Saveanu et al. 2003; Miles et al. 2005). Thus, based on this evidence coupled with the computational identification of RSL24D1 protein-protein interactions, we conclude that PeBoW members are RSL24D1-associated proteins in human cells.

**Table 1:**
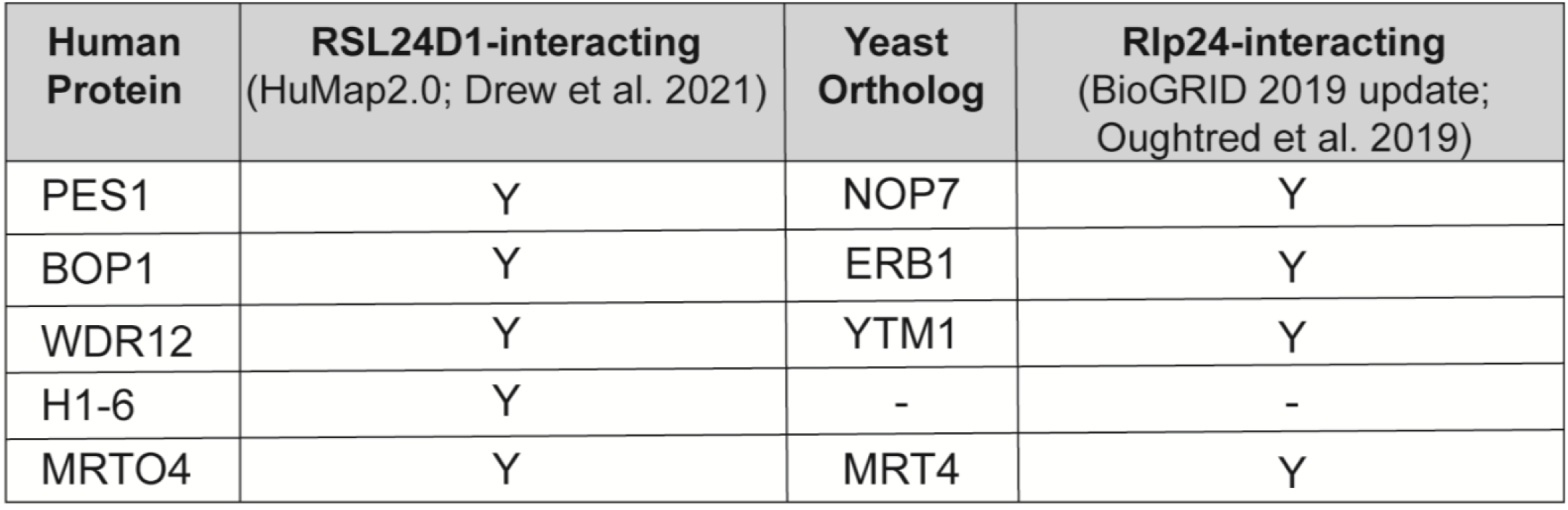

### RSL24D1 interacts with RNA polymerase I

Previously, Sorino and coworkers performed mass spectrometry of RPA194, the largest subunit of the RNAPI holoenzyme and identified RSL24D1 as a putative interacting partner (Sorino et al. 2020). Among RPA194 interacting proteins detected in their analysis, known RNAPI transcription factors including NOL11 and UBF (Panov et al. 2006; Freed et al. 2012), were identified along with, interestingly, WDR12 of the PeBoW complex. To validate whether RSL24D1 interacts with the RNAPI enzyme complex, we performed co-immunoprecipitation experiments followed by western blotting to test for physical association. Intriguingly, antibodies against RPA194, but not unconjugated beads, co-immunoprecipitated endogenous RSL24D1 from MCF10A whole cell extracts (Fig. 2A). Further, treatment of extracts with RNAse A did not abrogate this interaction (Fig. 2A). The reciprocal co-immunoprecipitation also revealed that RSL24D1 interacts with RPA194 (Fig. 2B). This finding led us to ask whether RSL24D1 could also influence ribosome biogenesis at the level of pre-rRNA transcription in addition to its regulatory role in the later pre-LSU processing steps. Since PeBoW components had previously been shown to be RSL24D1-interacting partners, and WDR12 had also been shown to co-purify with RPA194, we extended our investigation to include PeBoW complex members.

**Figure 2:**
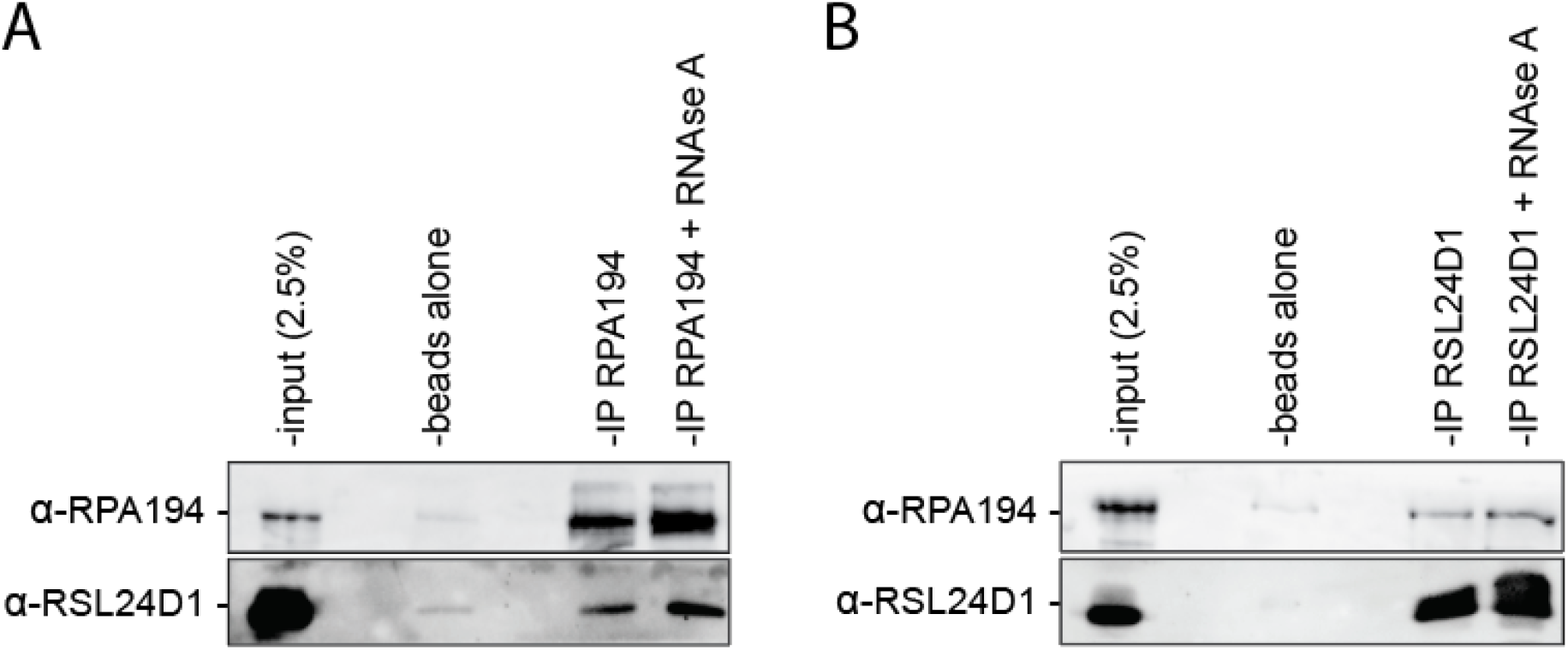
Co-immunoprecipitation analysis reveals an interaction between RSL24D1 and RNAPI. MCF10A whole cell extracts were immunoprecipitated with α-RPA194 **(A)** or α-RSL24D1 antibodies **(B)**. Conditions with/without RNase A treatment are shown. Input corresponds to 2.5% of the whole cell extract (WCE) used for immunoprecipitation. Unconjugated protein A beads were used as a negative control. RSL24D1 and RPA194 were detected by western blotting with specific antibodies.

### RSL24D1 and the PeBoW complex are required for RNAPI promoter activity

We first assessed pre-rRNA transcription by RNAPI using a dual-luciferase reporter assay. In this system, the pHrD-IRES-Luc plasmid, which contains the firefly luciferase gene under the control of the human rDNA promoter, is co-transfected with a control plasmid containing the Renilla luciferase gene under the control of a constitutively active promoter (Fig. 3A) (Ghoshal et al. 2004; Farley-Barnes et al. 2018). We assayed transcription in RSL24D1-, PES1-, BOP1- or WDR12-siRNA depleted MCF10A cells where siNT and siNOL11 were used as negative and positive controls, respectively. siRNAs targeting the Shwachman-Bodian-Diamond Syndrome (*SBDS*) transcript were employed as an additional negative control. SBDS is a pre-28S processing factor required for late-stage maturation of the large subunit of the ribosome and is not expected to be recruited for RNAPI transcription (Warren 2018). Remarkably, depletion of RSL24D1 or individual PeBoW members revealed reduced rDNA promoter activity and thus decreased pre-rRNA transcription (Fig. 3B). This result was further corroborated by in cellulo pulse labeling with the uridine analogue 5-ethynyl uridine (5-EU) and the analysis of its incorporation into nascent nucleolar rRNA through immunofluorescent staining as in (Bryant et al. 2022). Strikingly, cells depleted of RSL24D1 or PeBoW components exhibited a strong reduction in nucleolar rRNA biogenesis relative to siNT and siSBDS negative control cells (Fig. 3C, D), consistent with previous results following depletion of Ribi factors required for both pre-rRNA processing and transcription (Bryant et al. 2022). Taken together, these results indicate that RSL24D1 and PeBoW modulate rDNA transcription, linking LSU processing factors to RNAPI transcription regulation.

**Figure 3:**
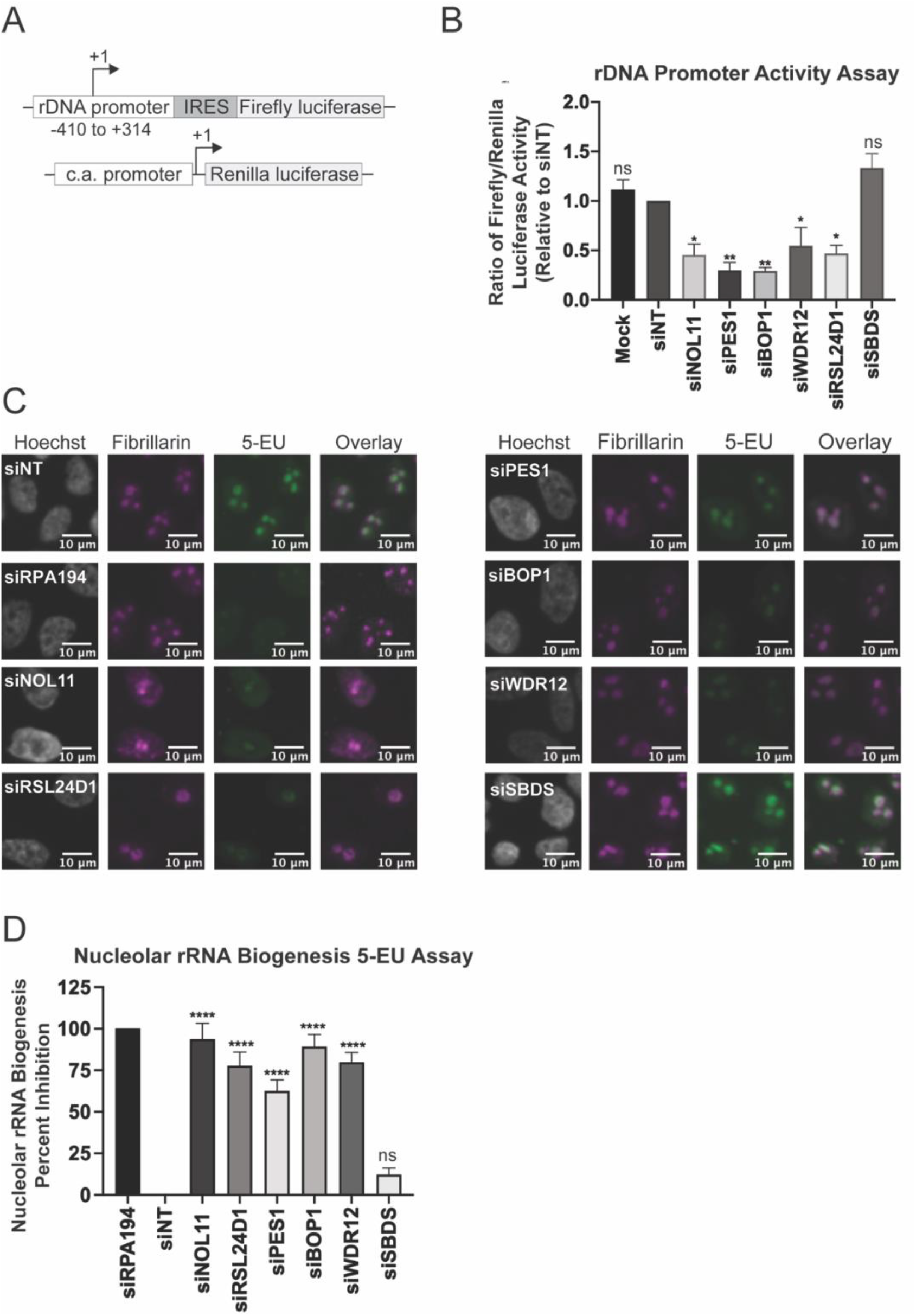
RSL24D1 and PeBoW complex members are required for optimal rDNA transcription. **(A)** Schematic of Firefly (pHrD-IRES-Luc) and Renilla luciferase plasmids; c.a. promoter: constitutively active promoter. **(B)** Depletion of RSL24D1 or PeBoW complex members leads to downregulated rDNA promoter activity. MCF10A cells treated with indicated siRNAs were transfected with reporter plasmids expressing Firefly and Renilla luciferase. Luminescence was quantified as a ratio in which Firefly gene expression, controlled by the human rDNA promoter, was normalized to Renilla gene expression, controlled by a constitutive promoter, and reported relative to siNT. Mock, siSBDS, and siNT were used as negative controls and siNOL11 was used as a positive control. Graph indicates mean ± SEM, n = 3 biological replicates. Data were analyzed by one-way ANOVA with Dunnett’s multiple comparisons test using GraphPad Prism where ** p ≤ 0.01, and * p ≤ 0.05; ns = not significant. **(C)** Nascent nucleolar RNA levels are reduced after siRNA depletion of RSL24D1 and individual PeBoW complex members by 5-EU visualization of nucleolar rRNA biogenesis. 72 h after siRNA transfection, MCF10A cells were supplemented with 1 mM 5-EU for 1 h before fixation. Fixed cells were stained for DNA (Hoechst) and the nucleolar protein fibrillarin (FBL), and click chemistry was performed to conjugate AF488 azide to labeled nascent RNA (5-EU). **(D)** Quantitation of the results in (C). Nucleolar rRNA biogenesis was quantified in MCF10A cells treated with the indicated siRNAs, where strongly reduced 5-EU signal corresponds to RNAPI inhibition (Bryant et al. 2022). The negative control non-targeting siRNA, siNT, (n = 16 wells per plate) was set at 0% inhibition and the positive control siRPA194 (n = 16 wells per plate) was set at 100% inhibition. Graph indicates mean ± SD, n = 3 biological replicates. Data were analyzed by one-way ANOVA with Dunnett’s multiple comparisons test in GraphPad Prism where **** p ≤ 0.0001; ns = not significant.

### RSL24D1 is required for sustained protein synthesis

Since RSL24D1 is required for pre-rRNA transcription and processing, we tested the extent to which RSL24D1 depletion impacts downstream processes, specifically global mRNA translation. We utilized a puromycin incorporation assay that labels nascent polypeptides as a proxy to monitor protein synthesis (Schmidt et al. 2009; Farley-Barnes et al. 2018). Treatment with 1 μM puromycin in RSL24D1-depleted MCF10A cells for 1 h followed by western blotting for incorporated puromycin shows significantly decreased protein synthesis relative to a siNT control (Fig. 4A, B). Mock (1 μM puromycin with no siRNA) and Mock at a half-dose of puromycin (0.5 μM) were used as negative controls whereas siNOL11 was a positive control. Our results connect RSL24D1’s role in pre-rRNA transcription and large subunit maturation to a downstream significant reduction in ribosome function upon its depletion.

**Figure 4:**
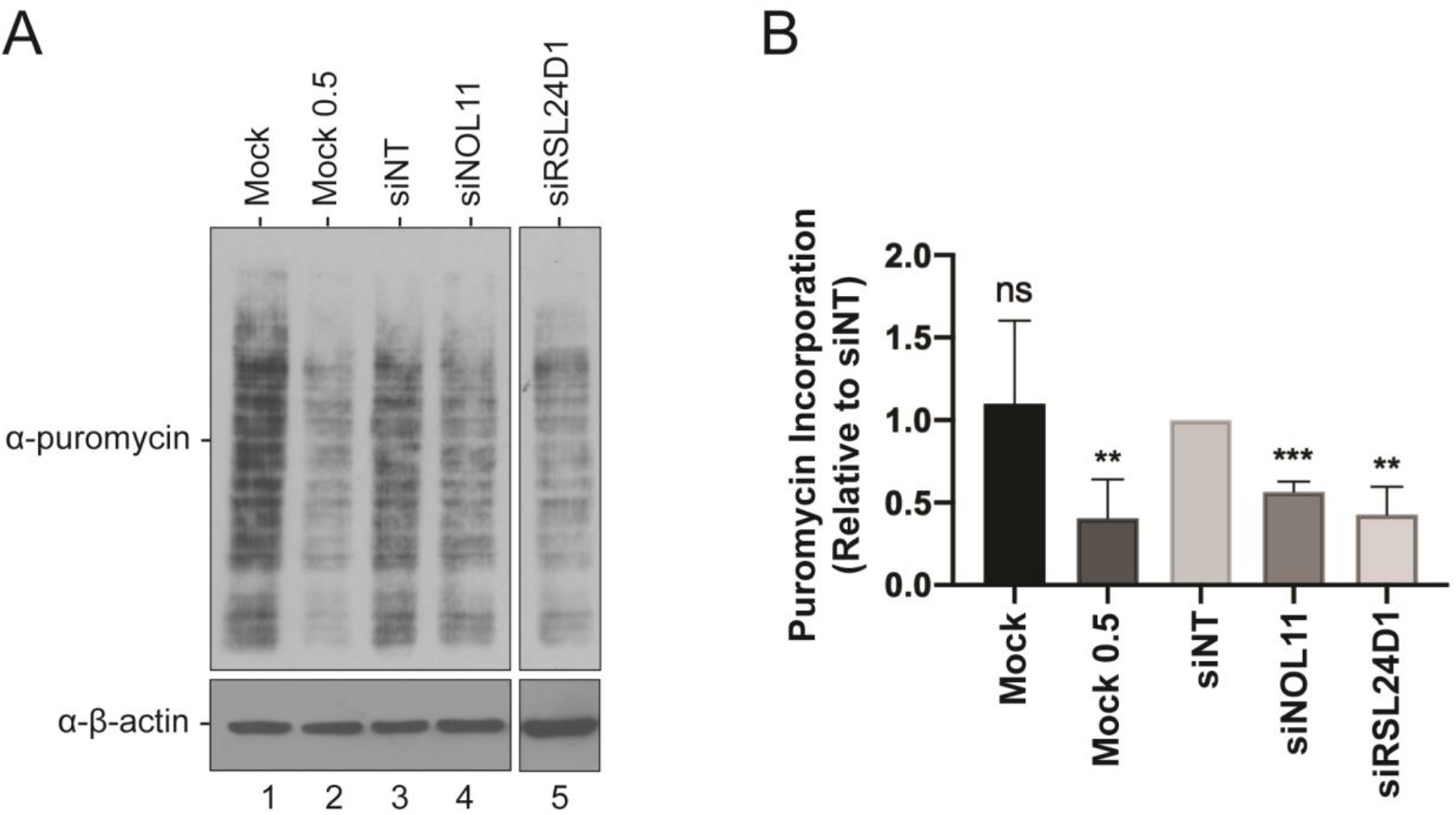
RSL24D1 depletion reduces global protein synthesis. **(A)** MCF10A cells were treated with 1 μM puromycin for 1 h following 72 h depletion with the indicated siRNAs. Harvested protein was analyzed by SDS-PAGE, followed by western blotting with an α-puromycin antibody. Mock (1 μM) and a non-targeting siRNA (siNT) were used as negative controls and siNOL11 was used as a positive control. Mock 0.5 μM indicates no siRNA and half the concentration of puromycin. β-actin was used as a loading control. **(B)** Quantitation of western blots from three biological replicates. ImageJ was used to quantify the differences in puromycin signal intensity, normalized to the β-actin signal intensity. Graph indicates mean ± SEM, n = 3 biological replicates. Data were analyzed by Student’s t-test followed by multiple testing p-value correction (two-stage linear step-up procedure of Benjamini, Krieger, and Yekutieli) using GraphPad Prism where *** p ≤ 0.001 and ** p ≤ 0.01; ns = not significant.

### RSL24D1 depletion induces the nucleolar stress response

Nucleolar stress denotes a key cellular response to drugs and environmental insults including impaired ribosome biogenesis due to RNAPI transcription repression (Rubbi and Milner 2003; Lindstrom et al. 2018). When disrupted, the nucleolus will often respond by activating the p53 pathway to initiate cell cycle arrest and apoptosis. Previously, PeBoW-complex members have been shown to induce the nucleolar stress response pathway when depleted (Grimm et al. 2006). Likewise, we tested whether nucleolar perturbations described in Fig. 1 upon RSL24D1 depletion were linked to concomitant p53 stabilization by western blotting for p53 levels in human RKO cells depleted of RSL24D1. Indeed, knockdown of RSL24D1 induced p53 accumulation (Fig. 5A). p53 stabilization following RSL24D1 knockdown was also observed in a recent genome-wide high-throughput screen conducted in A549 cells (Hannan et al. 2021). We further blotted for p53 transcriptional target gene p21 levels to orthogonally validate p53 activation and found these levels to be significantly increased relative to the mock and siNT negative controls (Fig. 5B). These findings show RSL24D1 depletion increases p53 levels, which may be attributed to repressed RNAPI transcription after loss of RSL24D1.

**Figure 5.**
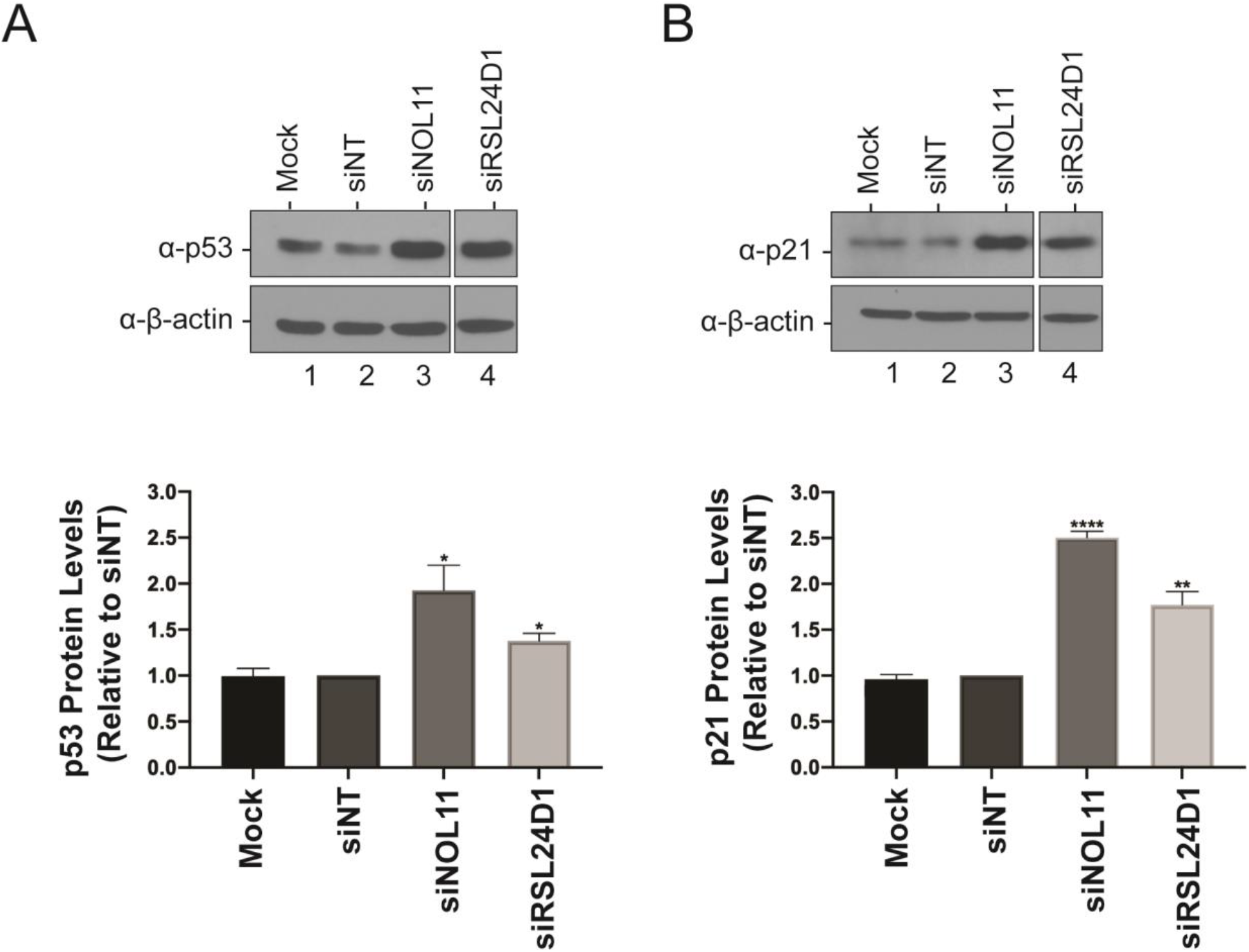
siRNA depletion of RSL24D1 causes the nucleolar stress response. **(A)** RSL24D1 depletion increases p53 protein levels. RKO cells were treated with indicated siRNAs for 72 h and total protein from whole cell extracts was harvested. Western blotting was performed for p53, and β-actin was used as a loading control. Non-targeting siRNA (siNT) and siNOL11 were used as negative and positive controls respectively, and data was normalized to siNT. Graph indicates mean ± SD, n = 3 biological replicates. Data were analyzed by Student’s t-test followed by multiple testing p-value correction (two-stage linear step-up procedure of Benjamini, Krieger, and Yekutieli) using GraphPad Prism, where * p < 0.05, ** p < 0.01, *** p < 0.001, **** p < 0.0001. **(B)** RSL24D1 depletion increases p21 protein levels. Experimental and statistical details are as above, except western blotting was performed for p21.

## Discussion

Here, we report the functional analysis of the putative ribosome biogenesis assembly factor RSL24D1. This protein was previously identified to be important for rRNA synthesis (Tafforeau et al. 2013; Durand et al. 2021), and consistent with this report, we demonstrate that RNAi mediated depletion of RSL24D1 specifically inhibits the late pre-rRNA processing steps required for 60S subunit formation. Moreover, repression of this pathway by RSL24D1 depletion leads to global reduction in mRNA translation. We show that disruption of these processes after knockdown of RSL24D1 is associated with a reduction in nucleolar number and concomitant activation of the nucleolar stress response pathway. Remarkably, we uncover an unexpected role for RSL24D1 and the PeBoW complex for the efficient production of pre-rRNA by RNAPI, the first step of ribosome biogenesis. We demonstrate that RSL24D1 is a positive regulator of rDNA promoter activity and directly interacts with RPA194, the largest subunit of the RNAPI enzyme complex. These results support a model that links LSU biogenesis factors to pre-rRNA transcription regulation.

In the present study, we provide strong evidence that LSU-associating assembly factors can also serve as dual-function factors in pre-rRNA transcription and processing. Several assembly factors associate with the nascent pre-rRNA transcript to facilitate RNAPI recruitment and initiation. A subgroup of these factors, the t-UTPs, assemble synchronously to promote rDNA transcription and stimulate the formation of the SSU processome for the processing of the pre-40S subunit (Gallagher et al. 2004; Krogan et al. 2004; Prieto and McStay 2007). The results presented here are substantiated by previous studies that have identified nucleolar proteins that regulate both ribosomal RNA transcription and pre-LSU processing, including splicing factor HTATSF1 and DNA repair protein FANCI (Corsini et al. 2018; Sondalle et al. 2019). We extend this framework to include the LSU factors RSL24D1 and PeBoW complex members (PES1-BOP1-WDR12) whose depletion leads to down-regulated rDNA promoter activity, strongly decreased nucleolar rRNA biogenesis consistent with roles in pre-rRNA processing and transcription, and impaired pre-LSU processing. Interestingly, PES1 has been identified to be an interacting partner of Upstream Binding Transcription Factor (UBTF) which is required for the recruitment of the pre-initiation complex along the rDNA promoter. Furthermore, all three PeBoW members interact with SIRT7, a positive regulator of RNAPI (Ford et al. 2006; Huttlin et al. 2017). Further studies investigating the crosstalk between these protein interactions and their influence on RNAPI transcription activation will be able to untangle the precise mechanism through which PeBoW complex members regulate this process.

While RSL24D1 gene mutations are not yet known to be implicated in the molecular pathogenesis of ribosomopathies, the diseases of making ribosomes, several were shown to be linked to mutations in genes encoding LSU biogenesis factors. For example, mutations in *SBDS* are associated with Schwachman-Diamond syndrome (Boocock et al. 2003) and defects in *RBM28* cause alopecia, neurological defects, and endocrinopathy (ANE) syndrome (Nousbeck et al. 2008; McCann et al. 2016; Bryant et al. 2021). Similarly, there are several diseases caused by mutations in ribosomal proteins of the LSU including Diamond-Blackfan anemia (*RPL5; RPL11; RPL27*) (Boria et al. 2010) and X-linked intellectual disability (*RPL10*) (Brooks et al. 2014). Intriguingly, RSL24D1 was recently reported to be required for murine embryonic stem cell proliferation (Durand et al. 2021). The prevalence of disorders arising from mutations in genes encoding ribosomal proteins and assembly factors of the large subunit highlight the importance of faithful subunit processing for the steady production of ribosomes to maintain cellular homeostasis.

## Materials and methods

### Publicly Available Expression Datasets

Genotype-Tissue Expression (GTEx) unmatched normal and The Cancer Genome Atlas (TCGA) matched normal and tumor expression datasets were obtained through the Xena platform (https://xena.ucsc.edu/) (Goldman et al. 2020). RNA-seq by Expectation-Maximization (RSEM) LOG2 fold expression levels for *RSL24D1* were subtracted from the mean of the overall normal and tumor tissues combined for graphical visualization. Expression values of 0 (not detected) were excluded from the graphs for ease of visualization for all tissues (normal = 26 excluded, tumor = 1 excluded).

### Cell culture and media

MCF10A cells (ATCC CRL-10317) were subcultured in Dulbecco’s modified Eagles’ medium/Nutrient mixture F-12 (Gibco 1130-032) containing horse serum (Gibco 16050), 10 μg/mL insulin (Sigma I1882), 0.5 μg/mL hydrocortisone (Sigma H0135), 100 ng/mL cholera toxin (Sigma C8052), and 20 ng/mL epidermal growth factor (Peprotech AF-100-15). RKO (ATCC CRL-2577) cells were grown in DMEM (Gibco 41965-062) with 10% fetal bovine serum (FBS, Gibco 10438026). Cell lines were maintained at 37 °C, in a humidified atmosphere with 5% CO_2_. For the high-throughput nucleolar number and 5-EU screens, 3,000 cells/well were reverse transfected into 384-well plates on day 0. For the dual-luciferase reporter assay, 75,000 cells/well were seeded into 12-well plates on day 0. For RNA or protein isolation, 100,000 cells/well were seeded into 6-well plates on day 0.

### RNAi

All siRNAs were purchased from Horizon Discovery Biosciences (Data S1). siGENOME SMARTpool siRNAs were used in the original screen (Farley-Barnes et al. 2018). For the 5-EU secondary screen validation of RSL24D1, PeBoW, and SBDS, ON-TARGETplus pools were used except for the NOL11 positive control which used the siGENOME SMARTpool siRNAs. For screen validation by deconvolution, the individual siONT set of four siRNAs that comprised the pool was used. The ON-TARGETplus pools were used in the remaining functional analysis. siRNA transfection was performed using Lipofectamine RNAiMAX Transfection Reagent (Invitrogen 13778150) as described in (Ogawa et al. 2021). For the high-throughput screens, cells were reverse transfected into 384-well plates on day 0. For other assays, cells were transfected 24 h after plating. For siONT deconvolution, an individual siRNA targeting RSL24D1 was considered validated if it produced a mean one-nucleolus percent effect greater than or equal to +3 SD above the siNT mean, using the siNT SD.

### 5-EU incorporation assay for nucleolar rRNA biogenesis

Following 72 h of siRNA depletion, MCF10A cells were treated with 1 mM 5-ethynyl uridine (5-EU; Click Chemistry Tools 1261) for 1 h to label nascent RNA as in (Bryant et al. 2022). Briefly, cells were washed with PBS, fixed with 1% paraformaldehyde (Electron Microscopy Sciences 15710-S) in PBS for 20 minutes, and permeabilized with 0.5% (vol/vol) Triton X-100 in PBS for 5 minutes. Cells were blocked with 10% (vol/vol) FBS (MilliporeSigma F0926) in PBS for 1 h at room temperature. Nucleoli were stained with 72B9 primary anti-fibrillarin antibody (Reimer et al. 1987), 1:250 for 2 h at room temperature, followed by secondary AlexaFluor 647 goat anti-mouse immunoglobulin G (1:1,000, Invitrogen A-21235) with Hoechst 33342 dye (1:3,000) for nuclei detection for 1 h at room temperature. 5-EU incorporation was visualized by performing the following click reaction in PBS: CuSO_4_:5H_2_O (0.5 mg/mL resuspended in water, Acros Organics 197730010), ascorbic acid (20 mg/mL freshly resuspended in water, Alfa Aesar A15613), and AF488 azide (0.5 μM resuspended in DMSO, Click Chemistry Tools 1275) for 30 min at room temperature. Cells were then soaked with Hoechst 33342 (1:3,000) for 30 min to dissociate excess azide dye. Cell images were acquired using IN Cell 2200 imaging system (GE Healthcare) and analysis was performed using a custom CellProfiler pipeline (Bryant et al. 2022).

### Dual-luciferase reporter assay for pre-rRNA transcription

Following 48 h of siRNA depletion, MCF10A cells were co-transfected with 1000 ng of pHrD-IRES-Luc (Ghoshal et al. 2004) and 0.1 ng of a Renilla internal control plasmid using Lipofectamine 3000 (Thermo Fisher Scientific L3000015). Twenty-four h after plasmid transfection, cells here harvested and luminescence was measured using the Dual-luciferase Reporter Assay System (Promega E1910) following the manufacturer’s instructions with a GloMax 20/20 luminometer (Promega). The ratio of pHrD-IRES-luciferase/Renilla activity was calculated to control for transfection efficiency and normalized to the non-targeting control.

### Northern blots

After 72 h of siRNA-mediated depletion in MCF10A cells, total cellular RNA was extracted with TRIzol (Life Technologies 5596018) according to the manufacturer’s protocol. For each sample, 4 μg of total RNA was resolved on a denaturing 1% agarose/1.25% formaldehyde gel using Tri/Tri buffer (Mansour and Pestov 2013) and transferred to a Hybond-XL membrane (GE Healthcare RPN 303S). Blots were hybridized to radiolabeled DNA oligonucleotide probes (P4: 5’-CGGGAACTCGGCCCGAGCCGGCTCTCTCTTTCCCTCTCCG-3’; 7SL: 5’-TGCTCCGTTTCCGACCTGGGCCGGTTCACCCCTCCTT-3’) and detected by phosphorimager as described previously (Pestov et al. 2008).

### qRT-PCR analysis

After 72 h of siRNA-mediated depletion, RNA was extracted using TRIzol (Life Technologies 5596018) as per the manufacturer’s instructions. cDNA preparation was performed using the iScript gDNA Clear cDNA synthesis Kit (Bio-Rad 1725035), and qPCR was performed using iTaq Universal SYBR Green Supermix (Bio-Rad 1725121). All A_260/230_ values were above 1.7 prior to cDNA synthesis. Amplification of the 7SL RNA was used as an internal control, and analysis was completed using the comparative C_T_ method (ΔΔC_T_). Bio-Rad PrimePCR Assay gene-specific primers were used to test *RSL24D1* mRNA levels (Bio-Rad 10025636; RSL24D1, qHsaCID0021318) and the following primers were used for the 7SL RNA forward: 5’ ATC GGG TGT CCG CAC TAA GTT-3’ and Reverse: 5’ – CAG CAC GGG AGT TTT GAC CT-3’ (Galiveti et al. 2010). Melt curves were performed for each sample to verify the amplification of a single product. Three biological replicates, each with three technical replicates, were measured.

### Western blots

Following 72 h of siRNA depletion, cells were collected and total protein was harvested and prepared as in (Farley-Barnes et al. 2018). 40 μg of total protein was separated by SDS-PAGE and transferred to a PVDF membrane (Bio-Rad 1620177). Membranes were blocked for 1 h in 5% milk in 1X PBST (or 5% BSA in 1X PBST for western blots of immunoprecipitations) and incubated overnight with the specified primary antibodies at 4 °C. Proteins were detected with the following antibodies: α-RSL24D1 (dilution 1:1,000; Santa Cruz sc-100840), α-RPA194 (dilution 1:1,000; Santa Cruz sc-48385), α-p53 (dilution 1:5,000; Santa Cruz sc-126), α-p21 (dilution 1:400, Santa Cruz sc-6246), α-puromycin (dilution 1:10,000; Kerafast EQ0001), and α-β-actin (dilution 1:30,000; Sigma Aldrich A1978). For the detection of the corresponding primary antibodies, either a 1:10,000 dilution of α-mouse-HRP conjugated antibody (GE Healthcare Life Sciences NA931) or a 1:2,000 dilution of α-mouse-HRP conjugated antibody (used for western blot of immunoprecipitations; Rockland, Mouse TrueBlot ULTRA, Ca. No. 18-8817-31) was used. The western blots were developed with enhanced chemiluminescence reagents (Thermo Scientific 34096). Images were acquired by digital imaging using the Bio-Rad ChemiDoc Imaging System. Images were quantified with ImageJ software.

### Puromycin labeling assay

Global protein translation was assessed by treating cells with puromycin to label nascent polypeptide chains as in (Schmidt et al. 2009). Total cellular extracts were then analyzed by western blot using an anti-puromycin antibody (Kerafast EQ0001) at a 1:10,000 dilution. Quantitation was performed using ImageJ.

### Co-immunoprecipitation

Protein A agarose beads (Cell Signaling Technologies 9863S) were washed and incubated in NET150 buffer (20 mM Tris HCl ph. 7.5, 150 mM NaCl, 0.05% NP-40) with 20 μg of the indicated antibodies overnight at 4 °C with nutation. Harvested MCF10A cells were washed with PBS and incubated on ice for 10 min in NET150 buffer (with the addition of 1X protease inhibitors and 4 mM NEM). Total cell extracts were obtained by sonication and cleared by centrifugation at 16,000 g for 10 min at 4 °C after lysis. Supernatants were incubated with either antibody-bound or unconjugated Protein A beads for 2 h at 4 °C with nutation. RNase A treated extracts were treated with 20 μg/mL RNase A (AMRESCO E866). After beads were washed five times with NET150, immunocomplexes were eluted in 2X Laemmli buffer and resolved on a 12% acrylamide gel. Western blotting was performed as described above.

### Statistical analyses

All statistical analyses were performed in GraphPad Prism 8.2.1 (Graphpad Software) using the tests described in the figure legends.

## Acknowledgements

We thank current and former members of the Baserga laboratory for helpful discussions and insights in the preparation of this manuscript. This work was supported by the following grants from the National Institutes of Health (NIH): 1R35GM131687 (to S.J.B.), 1F31DE030332 (to M.A.M.), and T32GM007223 (to C.J.B., M.A.M., and S.J.B.).

