## Supplementary Data and Figures for "Human pre-60S assembly factors link rRNA transcription to pre-rRNA processing"

**Data S1: Horizon Discovery siRNAs used in this study**

| siRNA target | Catalog Number |
| --- | --- |
| siNT (ON-TARGETplus SMARTpool) | D-001810-10-20 |
| siNOL11 (ON-TARGETplus SMARTpool) | M-016695-01-0005 |
| siRSL24D1 (ON-TARGETplus SMARTpool) | L-013170-02-0005 |
| siRSL24D1-1 (ON-TARGETplus - Individual) | J-013170-17-0002 |
| siRSL24D1-2 (ON-TARGETplus - Individual) | J-013170-18-0002 |
| siRSL24D1-3 (ON-TARGETplus - Individual) | J-013170-19-0002 |
| siRSL24D1-4 (ON-TARGETplus - Individual) | J-013170-20-0002 |
| siRPA194 (ON-TARGETplus SMARTpool) | L-013983-01-0005 |
| siPES1 (ON-TARGETplus SMARTpool) | L-009542-00-0005 |
| siBOP1 (ON-TARGETplus SMARTpool) | L-014065-01-0005 |
| siWDR12 (ON-TARGETplus SMARTpool) | L-012972-00-0005 |
| siSBDS (ON-TARGETplus SMARTpool) | L-019217-00-0005 |

siRSL24D1	siRSL24D1	siRSL24D1	siRSL24D1

siNT	siNOL11	individual #1	individual #2	individual #3	individual #4

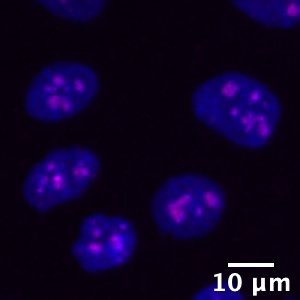

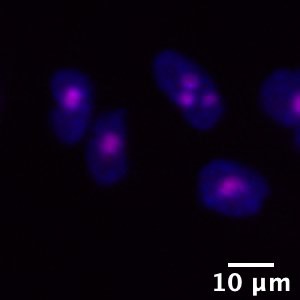

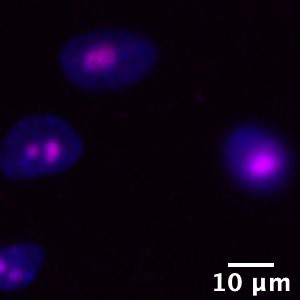

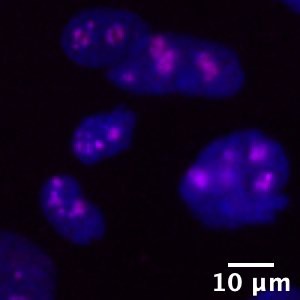

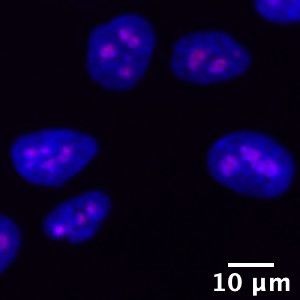

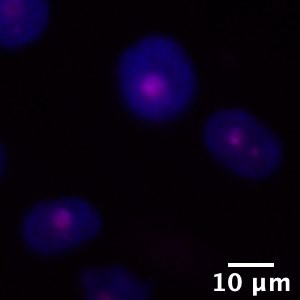

**Fig. S1.** **Representative images that show that two of four RSL24D1 siRNAs (#2 and #3) decrease nucleolar number in MCF10A cells.** Nuclei are stained with Hoechst (blue) and nucleoli stained with an anti-fibrillarin antibody (pink) in MCF10A cells treated with siON-TARGETplus siRNA reagents. Non-targeting siRNA (siNT) was used as a negative control, siNOL11 was used as a positive control, and four separate siRSL24D1 treatments are shown from each individual siRNA that compromises the siRSL24D1 ON-TARGETplus pool.

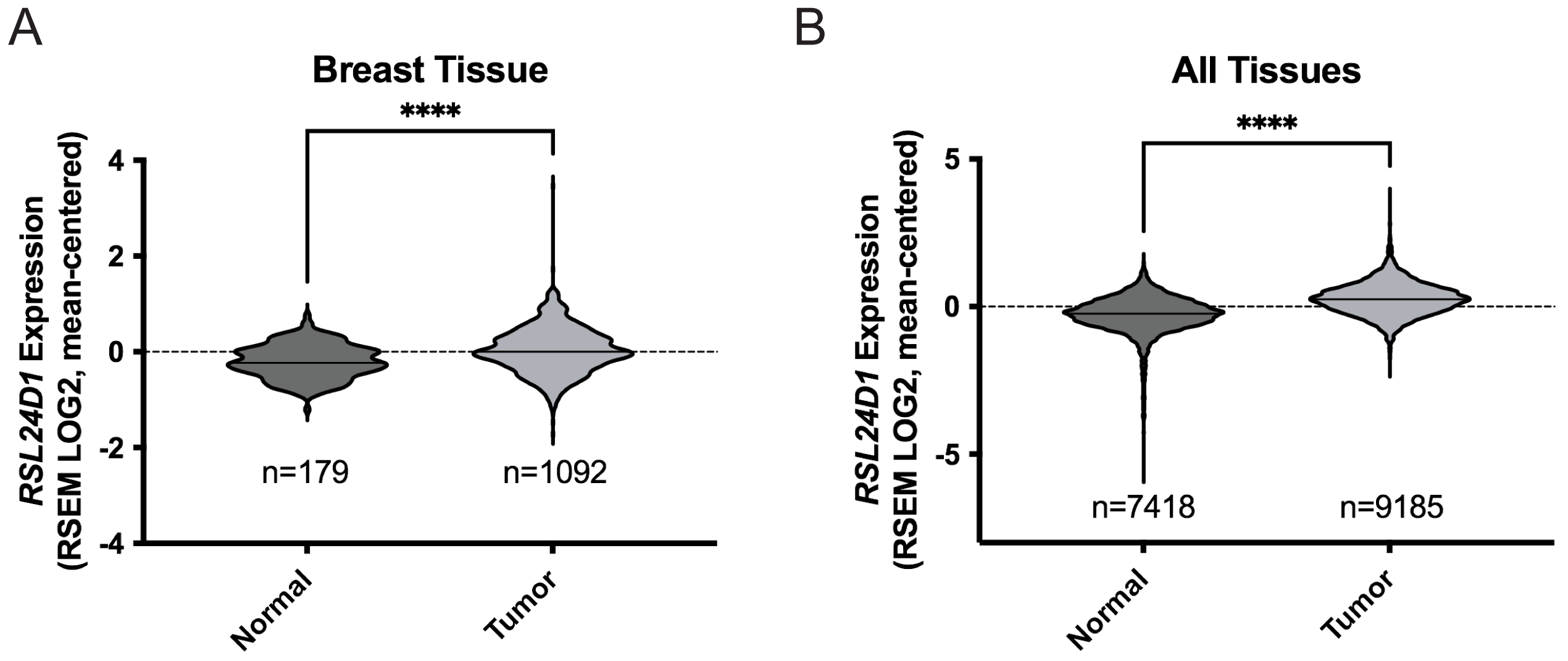

**Fig. S2. *RSL24D1* has increased expression levels in all cancers, including breast cancer.**

Violin plots of Genotype-Tissue Expression (GTEx) unmatched normal and The Cancer Genome Atlas (TCGA) matched normal and tumor RNA-seq by Expectation-Maximization (RSEM) LOG2 fold expression levels for *RSL24D1* subtracted from the mean. **(A)** *RSL24D1* expression in normal and tumor breast tissue. **(B)** *RSL24D1* expression in all normal and tumor tissues. Dashed line (set at 0) indicates mean of entire dataset for both normal and tumor expression, black lines indicate mean of individual normal or tumor expression dataset. Data were analyzed by Student’s t-test using GraphPad Prism where **** p ≤ 0.0001.
